# Induction of Rare Conformation of Oligosaccharide by Binding to Calcium-dependent Bacterial Lectin: X-ray Crystallography and Modelling Study

**DOI:** 10.1101/612689

**Authors:** Martin Lepsik, Roman Sommer, Sakonwan Kuhaudomlarp, Mickaёl Lelimousin, Emanuele Paci, Annabelle Varrot, Alexander Titz, Anne Imberty

**Author notes:** Corresponding author. Martin Lepsik & Anne Imberty.

## Abstract

Pathogenic micro-organisms utilize protein receptors in adhesion to host tissues, a process that in some cases relies on the interaction between lectin and human glycoconjugates. Oligosaccharide epitopes are recognized through their three-dimensional structure and their flexibility is a key issue in specificity. In this paper, we analyse by X-ray crystallography the structures of the lectin LecB from two strains of *Pseudomonas aeruginosa* in complex with Lewis x oligosaccharide present on cell surfaces of human tissues. An unusual conformation of the glycan was observed in all binding sites with a non-canonical *syn* orientation of the *N*-acetyl group of *N*-acetyl-glucosamine. A PDB-wide search revealed that such an orientation occurs only in 2% of protein/carbohydrate complexes. Theoretical chemistry calculations showed that the observed conformation is unstable in solution but stabilised by the lectin. A reliable description of LecB/Lewis x complex by force field-based methods had proven as especially challenging due to the special feature of the binding site, two closely apposed Ca^2+^ ions which induce strong charge delocalisation. By comparing various force-field parametrisations, we design general protocols which will be useful in near future for designing carbohydrate-based ligands (glycodrugs) against other calcium-dependent protein receptors.

## INTRODUCTION

*Pseudomonas aeruginosa* is a gram-negative bacterium, which acts as an opportunistic pathogen responsible for severe bronchopulmonary infections notably in cystic fibrosis patients. *P. aeruginosa* disposes of many soluble virulence factors, including two lectins, LecA (PA-IL) and LecB (PA-IIL), with specificity for galactose (Gal) and fucose (Fuc), respectively [1, 2]. Structurally, LecB is a homotetramer with four binding sites, each of which contains two Ca^2+^ atoms [3] that mediate the binding of Fuc accompanied by a strong charge delocalisation [4]. The high affinity to Fuc results in strong attachment to branched fucosylated oligosaccharides (Fig. 1A and 1B) present in the human blood group epitopes such as the AB(H) and Lewis series (Fig. 1C) [5].

**Figure 1.**
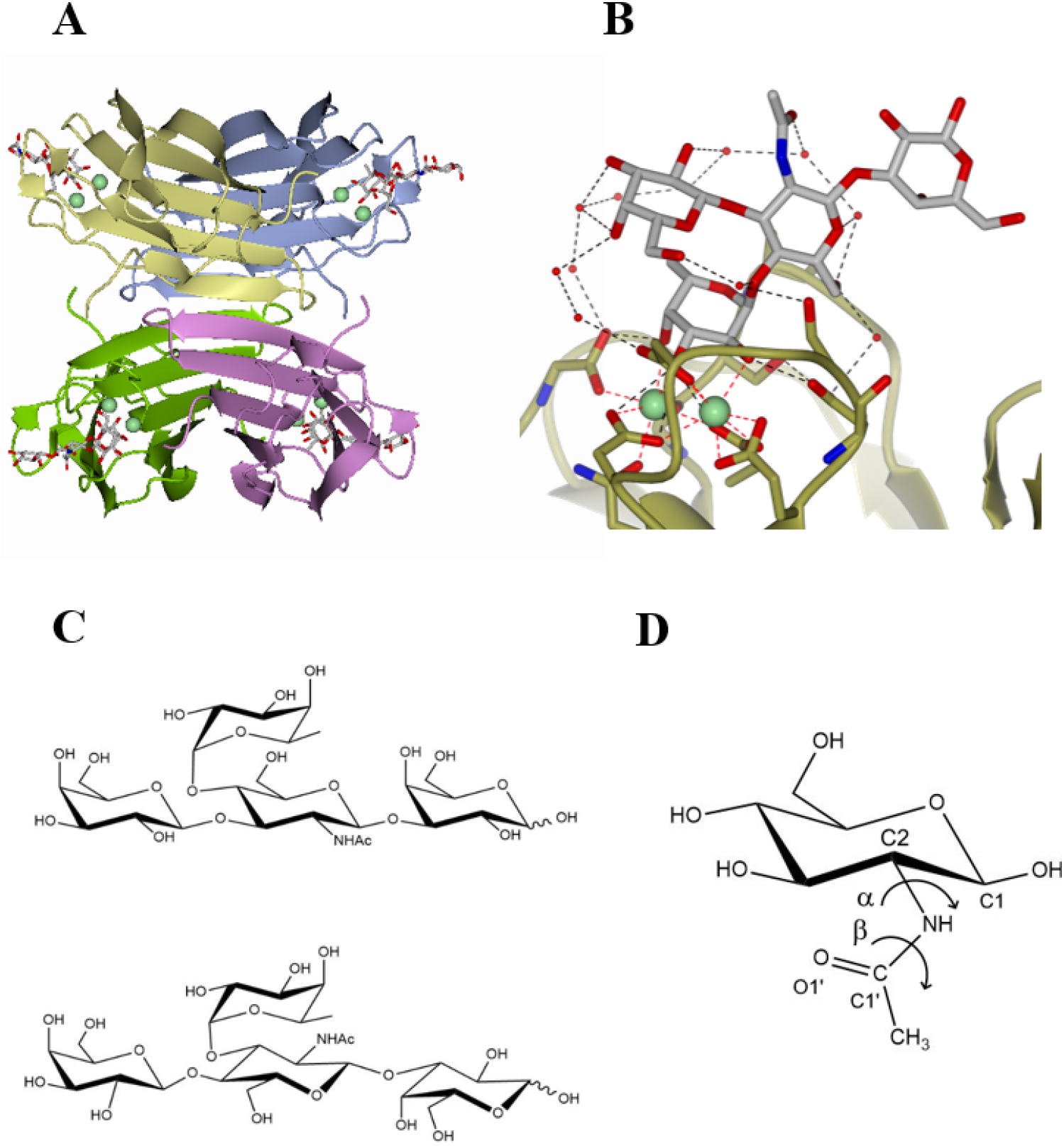
Structure of LecB and fucosylated ligands. **A** Overall view of LecB_PA14_ tetramer (coloured cartoon) with two calcium ions (green spheres) and one Le^a^ tetrasaccharide bound (sticks) in each binding site (PDB 5A6Z)[6]. **B** Zoom in the binding site displaying the orientation of the *N*-acetyl group in the solvent. **C** The chemical structures of Le^a^ (top) and Le^x^ (bottom) tetrasaccharides. **D** The chemical structure of d-GlcNAc with definitions of α- and β-dihedrals [7].

Among human epitopes, LecB demonstrated higher affinity for Lewis a oligosaccharide (Le^a^: Fuc α(1-4) [Gal β(1-3)] GlcNAc) which was rationalised by X-ray structure of the complex [8]. In the binding site, Fuc coordinates the two calcium ions via its hydroxyl groups, which moreover also act as hydrogen bond donors to acidic amino acids of the binding site. Furthermore, *N*-acetyl-glucosamine (GlcNAc) creates an additional hydrogen bond with the protein surface. Different *P. aeruginosa* strains have been investigated [6, 9], since the classical PAO1 laboratory strain is not the only to be involved in clinical infection. LecB from strains PA14 (LecB_PA14_) and PAO1 (LecB_PAO1_) differ by 13% in sequences, but both bind to Le^a^ with high affinity as determined by titration microcalorimetry (*K*_d_ of 78 nM and 170 nM, respectively) [6]. Binding to Lewis x epitope (Le^x^: Fuc α(1-3) [Gal β(1-4)] GlcNAc) is almost as efficient as to Le^a^ with affinities to LecB_PA14_ and LecB_PAO1_ of 90 and 400 nM, respectively) [6, 8]. This was unexpected since the orientation of GlcNAc in Le^x^ is different from that in Le^a^ and the *N*-acetyl was predicted to create steric conflict with the protein surface [3].

The Lewis epitopes are the targets for viral receptors, such as those of noroviruses, and for a number of bacterial lectins [5, 10]. These oligosaccharides are usually described as rigid due to their branched structure, that results in stacking between Fuc and Gal rings, steric hindrance of the *N*-acetyl group of *N*-acetylglucosamine (GlcNAc) and the presence of an unconventional CH⋯O hydrogen bond [11-13]. The conformation in solution of Lewis oligosaccharides is referred to as the “closed” conformation, which is the one recognized by most protein receptors. However, a class of lectins from fungi and bacteria has been reported to induce a conformation change in Le^x^ and a series of “open” conformations were observed by X-ray crystallography and rationalised by molecular dynamics (MD) calculations [14]. Characterising the interaction of bacterial lectins with their natural ligands is of interest for the development of anti-infective compounds that are able to inhibit the adhesion of bacteria or the formation of biofilm [15]. For example, the structure of LecB complexed with Le^a^ was used for the design of fucose derived glycomimetics with aromatic aglycone mimicking the GlcNAc ring [16] and mannose/fucose derived glycomimetics carrying sulphonamide substituents [17, 18].

We describe here the crystal structures of LecB from PAO1 and PA14 strains in complex with Le^x^ and we analyze the conformation of GlcNAc (Fig. 1D) by quantum mechanical (QM) and molecular dynamics (MD) calculations. The presence of bridging calcium ions and water molecules in the binding site necessitates special care for force-field parameterization as previously demonstrated [19, 20] and several approaches are compared. The derived method will be of general interest in the future for design of active compounds against calcium-containing lectins from pathogens.

## RESULTS

### Crystal structures of LecB complexed with Le^x^

LecB_PA14_ and LecB_PAO1_ were co-crystallised with the Le^x^ tetrasaccharide, resulting in crystals in P21 space group diffracting to 1.6 Å and 1.8 Å, respectively. The structures were solved by molecular replacement and details are listed in Table S1. Clear electron density was observed for the whole oligosaccharides in all four binding sites of both structures, except for the reducing-end Gal in LecB_PAO1_ (Fig. S1). Consistent with previous findings, both structures are homotetramers with four binding sites, each of which contains two Ca^2+^ atoms[3, 6]. In the same manner as for complexes with Fuc [3] or Le^a^ [6, 8], the calcium atoms mediate the binding of Fuc residue in Le^x^ via three hydroxyl oxygens (O2, O3 and O4). Additionally, the three hydroxyl groups of Fuc can form several direct hydrogen bonds with amino acid residues within the binding site (N21:O, D96:OD1, D99:OD2, D101:D2 and G114*:OXT). The recognition of Fuc residue by LecB_PA14_ and LecB_PAO1_ are in complete agreement with previous observations [3, 6].

In order to rationalise the recognition of Le^x^ by the LecB variants (LecB_PA14_ and LecB_PAO1_) in relation to the amino acid mutations in their binding site (S97/A23 vs G97/S23, respectively), detailed comparison of their complexes with Le^x^ was performed. The serine at position 97 in LecB_PA14_ provides a hydrophilic environment and organises waters W1, W2, and W3 to form several hydrogen-bond bridges between the protein and the carbohydrate (Fig. 2A). As a result of the S97G variation in LecB_PAO1_, W2 and W3 are absent in LecB_PAO1_-Le^x^ complex and the water-coordinated interaction is replaced by a direct hydrogen bond between the *N*-acetyl of the GlcNAc and the D96 side chain Fig. 2B). The W1 interactions with the carbohydrates are retained in both proteins. From the protein side, W1 is held by an H-bond from the N98:N backbone.

**Figure 2.**
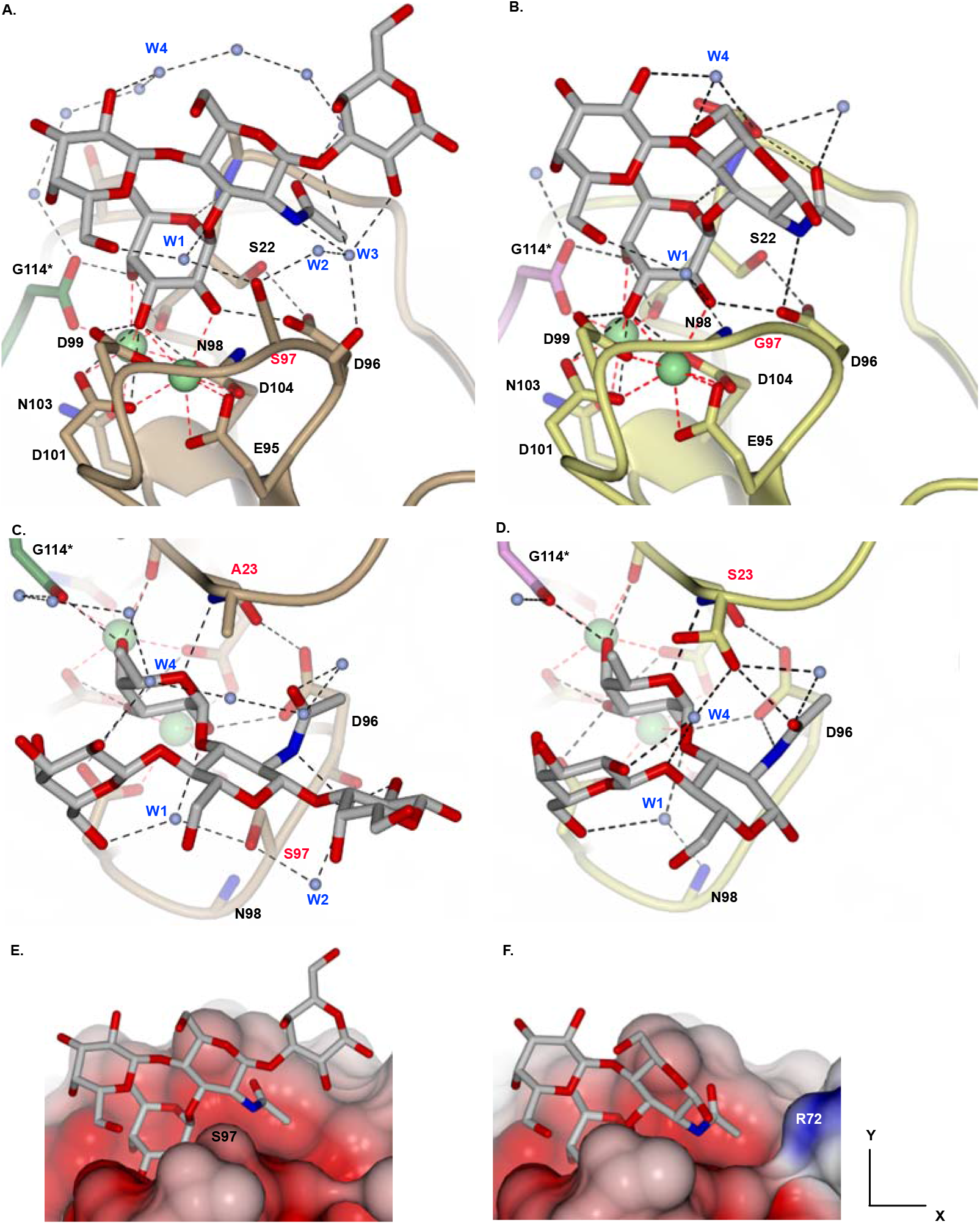
Comparison of crystallographic structures of Le^x^ in complex with LecB_PA14_ **(A, C)** and LecB_PAO1_ **(B, D)**. The views in **C, D** correspond to a 90° clockwise rotation of **A, B**, respectively, around the x axis. Ca^2+^ atoms are represented by green sphere. The protein backbones of LecB_PA14_ and LecB_PAO1_ are coloured in light brown and yellow, respectively. G114* is the N-ter residue of the adjacent monomer and is coloured in green (in LecB_P_A14) or pink (in LecB_P_AO1). Water molecules are shown as ice-blue spheres. Le^x^ is coloured in light grey (note that in LecB_PAO1_, the reducing galactose residue could not be modelled due to poor electron density; Fig. S1). **E** and **F** are surface representation of **A** and **B**, respectively. The surfaces are coloured by electrostatic potential (red; negative, blue; positive). Note that R72 is present in both LecB variants, but the side chain of R72 was only modelled in LecB_PAO1_, thus is only visible in **F**. The amino acid variants are labelled in red. The figure was rendered in CCP4MG.

The A23 variant in LecB_PA14_ constitutes (together with T45) a hydrophobic pocket which accepts the Fuc:C6 methyl (Fig. 2C), as described previously for LecB_PA14_-Le^a^ [6]. Further toward the bulk solvent, a water network links the protein with the carbohydrate (Fig. 2C). W4, present in this water network forms a direct contact with the non-reducing Gal (Fig. 2A, C). In LecB_PAO1_, S23 was found in two alternate conformations, one of which provides a direct H-bond to GlcNAc:O1’ (Fig. 2D). The presence of the S23 polar side chain seems to disrupt the water network described in LecB_PA14_ complex (Fig. 2D).

Due to the amino acid variations, some differences are observed in the interaction between Le^x^ and the two LecB variants. While the Fuc binding is virtually unchanged, the Φ, Ψ torsion angles of the *α*Fuc1-3GlcNAc glycosidic linkage in chains A and C differ by up to 16.5° and 11.5°, between LecB_PAO1_ and LecB_PA14_ (Table S2), which results in a slight change in the interactions. However, in chain D, these torsions are very similar, with changes of 0.9 and 0.5° (Table S2). Main observed differences are in the interaction between GlcNAc and the protein surface. In LecB_PAO1_, the NH of GlcNAc is close enough to the Asp96 side chain to form a direct hydrogen bond (Fig. 2B, D). In LecB_PA14_, the GlcNAc ring is pushed slightly away and instead of a direct hydrogen bond, a water bridge via W3 is observed between NH of GlcNAc and the backbone oxygen of Asp96 (Fig. 2A, C). In addition, the S97G mutation in LecA_PAO1_ provides a hydrophobic patch, enabling a closer contact of the GlcNAc ring with the protein surface, which is not observed in LecB_PA14_ due to the polar OH group of S97 side chain (Fig. 2E and F).

In the crystal structures of both LecB variants interacting with Le^x^, an unusual conformation is observed for the *N*-acetyl group of all four Le^x^ molecules of the asymmetric unit. In all cases, the C2-N bond adopts a non-canonical *syn* orientation of the α dihedral angle (Fig. 1D, Table 1). A detail of the electron density is displayed in Fig. S1. The β torsion angles always remain in the classical *trans* conformation (Fig. 1D, Table 1). In all the binding sites, the unusual conformation of the *N*-acetyl brings the bulky acetyl group away from the protein surface. The NH moiety points in the direction of the acidic group of Asp96, establishing a direct hydrogen bond in LecB_PAO1_, and a water bridged one in LecB_PA14_ (Table 1). The presence of S97 in the LecB_PA14_/Le^x^ complex and accompanying hydration results therefore in slightly different position and hydrogen bond network from the LecB_PAO1_/Le^x^ complex.

**Table 1.** Geometric characteristics of *N*-acetyl of Le^x^ in the crystal structures of complexes with LecB_PA14_ and LecB_PAO1_. α and β torsion angles (°) are defined as α (H2 – C2 – N – H), β (H – N – C1’– O1’) and d1 (Å) is the distance between GlcNAc:N and D96:OD1.

| chain | $\alpha$ | | $\beta$ | | d1 | |
| --- | --- | --- | --- | --- | --- | --- |
| | $\text{LecB}_{\text{PA14}}$ | $\text{LecB}_{\text{PAO1}}$ | $\text{LecB}_{\text{PA14}}$ | $\text{LecB}_{\text{PAO1}}$ | $\text{LecB}_{\text{PA14}}$ | $\text{LecB}_{\text{PAO1}}$ |
| A | -6.8 | -8.3 | 172.4 | 177.1 | 3.7 | 3.0 |
| B | -9.0 | -11.7 | 174.1 | 169.5 | 3.6 | 3.1 |
| C | -6.9 | -13.3 | 171.7 | 161.3 | 3.5 | 3.1 |
| D | -1.8 | 12.4 | 169.1 | 149.7 | 3.5 | 3.4 |

### Analysis of GlcNAc *N*-acetyl group in the Protein Data Bank

In order to evaluate the frequency of the rare non-canonical orientation of *N*-acetyl, the GlcNAc residues were analyse in the Protein Data Bank (PDB). Such a study was previously performed for glycoproteins and protein/glycan complexes [21], but with no quality check on the electron density. The search (as of March 17, 2019) yielded altogether 381 conformers, with 338 in the canonical α-*anti*/β-*trans* conformation i.e. 88.7 % of occurrences with dihedrals ranging from 139° to 209° (-151°) for α and 140° to 210° (-150°) for β (Fig. 3).

**Figure 3.**
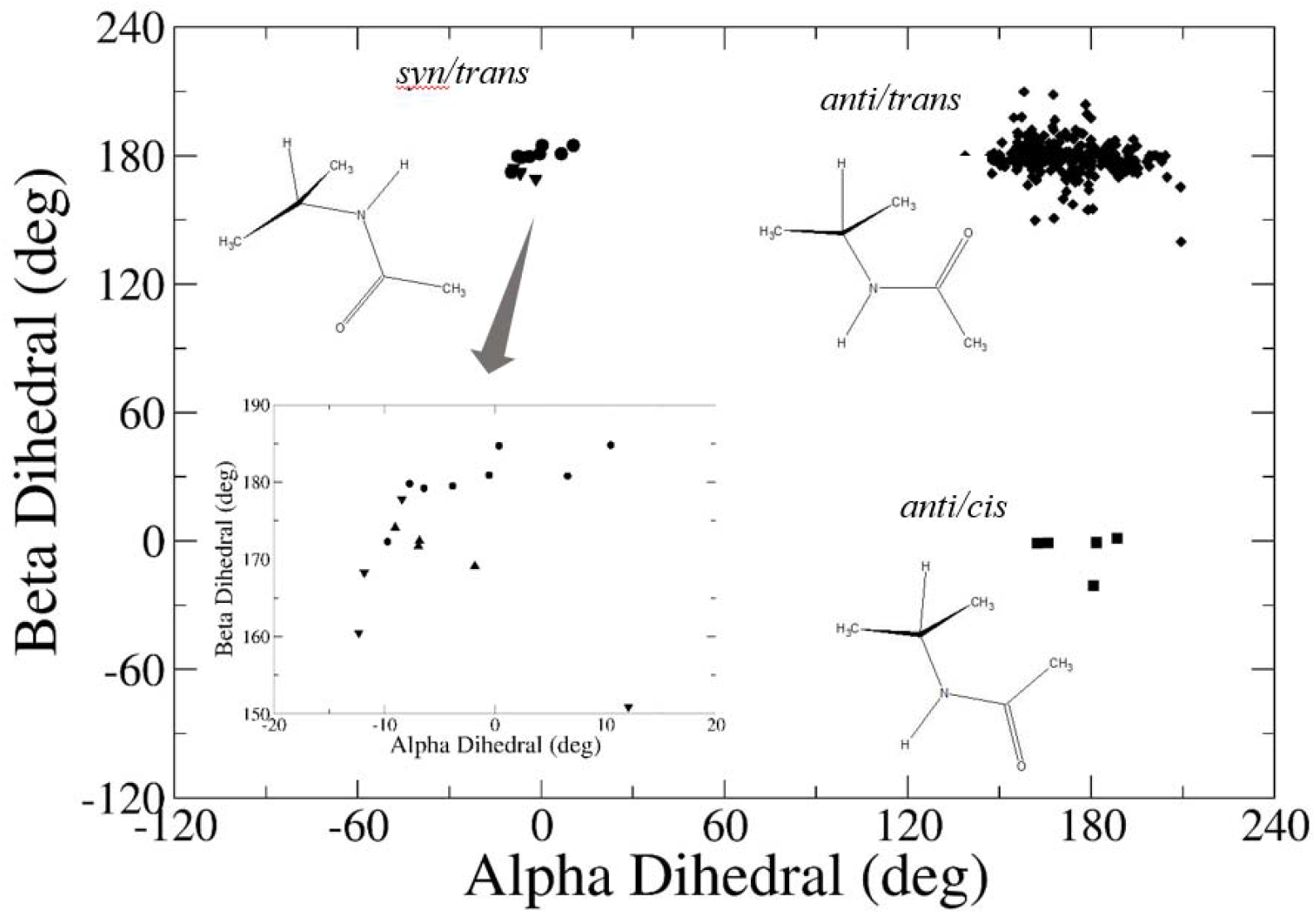
Analysis of *N*-acetyl α and β dihedral angles in β-d-GlcNAc in protein/carbohydrate complexes in the PDB (each point represents one monosaccharide in a PDB structure). The relative orientation of the functional groups in the minor populations is shown schematically. Inset: Zoom in the α-*syn*/β-*trans* region (crystal structures determined in this work; LecB_PA14_/Le^x^, triangle up; LecB_PAO1_/Le^x^, triangle down).

The other observed structures can be assigned to two non-canonical conformations, *syn/trans* or *anti/cis,* and one outlier *syn/cis.* These minor populations were further analysed by direct examination of the electron density maps using Coot [22] and the structure which were not supported by the electron density were discarded. For *syn/trans* torsion combination, 16 cases were confirmed including 8 from this work, i.e. 4 *%* occurrence with α ranging from -10° to 11° and β from 172° to 185° (-175°) (Fig. 3). For *anti/cis*, only six case were retained, i.e. 1.6% occurrence with α ranging from 165° to 188° (-172°) and β from -21° to 1° (Fig. 3). The *syn/cis* outlier was not supported by the electron density and as such it was discarded.

Analysis of the PDB complemented with examination of the electron density maps confirmed eight cases plus another eight observed here crystallographically (i.e. 4% of the torsions studied) of the *N*-acetyl α dihedral in *syn* conformation

### Conformational behaviour of the *N*-Acetyl group in Le^x^

The Le^x^ trisaccharide conformational behaviour was previously studied by us [14] by molecular dynamics using the GLYCAM06-j force field [23] in explicit water (TIP3P model) [24]. Several independent trajectories of 1 to 10 μs were produced, starting from the Le^x^ solution shape with the *N*-acetyl group in the canonical conformation. The behaviour of the *N*-acetyl group was analysed for the present work and no variations were observed for the *N*-acetyl group in these trajectories (data not shown).

In the present work, a 1.4 μs MD simulation of Le^x^ tetrasaccharide in TIP3P water was started from the α-*syn* conformation from the LecB_PA14_/Le^x^ complex crystal structure (chain A, see above). We observed a rapid conversion within 50 ns to α-*anti* and its persistence until the end of the simulation (Fig. 4A). The β dihedral stayed in all the cases in *trans* in range from 147° to 221° (-139°) (Fig. 4A). Therefore, the MD of the Le^x^ in aqueous solution confirms the stability of the major *anti/trans* conformation observed in most of the protein/carbohydrate complexes in the PDB. However, the determination of the rotational barrier is necessary in order to obtain trajectories with relevant occupancy of the different states.

**Figure 4.**
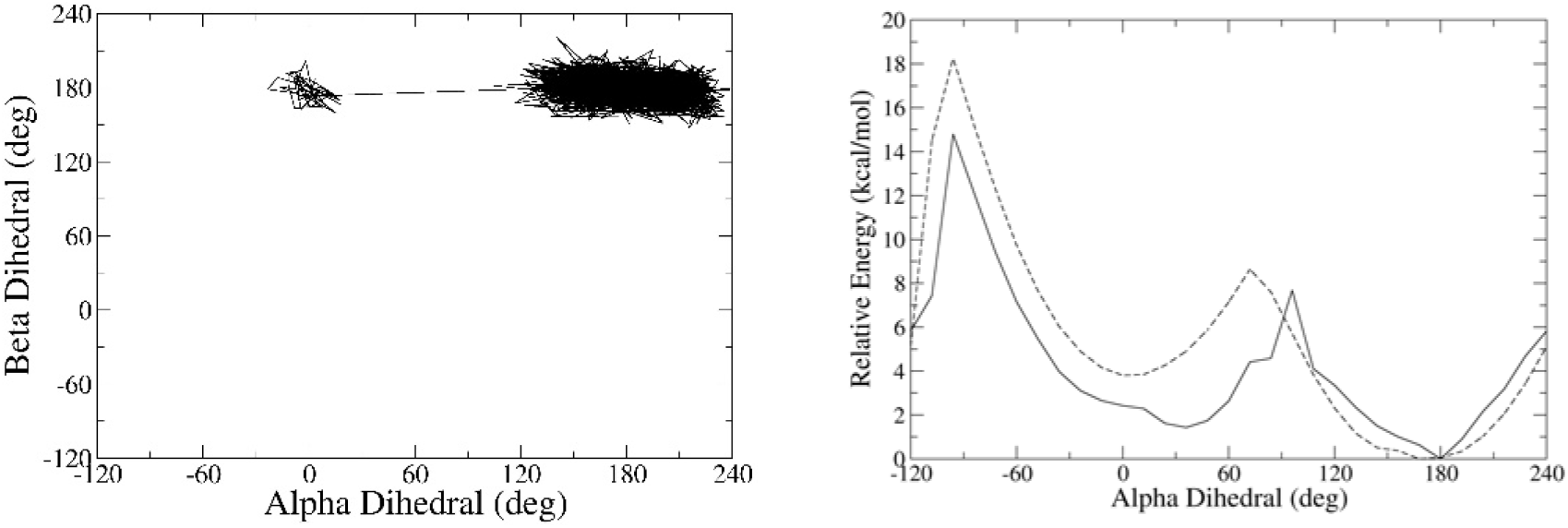
Conformational stability of the *N*-acetyl α dihedral of Le^x^ tetrasaccharide. **A**. MD/explicit solvent. *Syn/trans* (0/180°) changed to *anti/trans* (180/180°) after 50 ns and stayed until 1.4 μs. **B**. The relaxed rotational scan of the *N*-acetyl α dihedral in Le^x^ using quantum mechanics (solid line) and molecular mechanics (dashed line) calculations in implicit solvent (for details, see text).

The relaxed rotational scans of the dihedral in Le^x^ were calculated at two levels (Fig. 4B). First, for reference, we used quantum mechanics (QM; density functional theory with empirical dispersion (DFT-D3)[25] with B-LYP functional and double-ζ basis set (DZVP) [26] coupled with implicit solvent COSMO model [27] in Turbomole [28].Second, for comparison with the MD simulations, we used molecular mechanics (MM; GLYCAM06-j)[23] coupled with generalised Born (GB) implicit solvent using the *igb5* parametrisation [29] in AMBER18 [30]. Overall, both levels of theory give similar shapes of the curves. The locations and relative stabilities of the minima in Le^x^ mirror the populations found in the PDB (c.f. Fig. 3) and MD (Fig. 4A). The global minimum is at α-*anti* (180°) and the local minimum at α-*syn* (0°; relative energy to the minimum of the method is 3.8 kcal/mol for MM and 36° and 1.4 kcal/mol for QM). Barriers to rotation of the *N*-acetyl α dihedral from *syn* to *anti* are asymmetric. The high barriers at -96° of 14.5 kcal/mol (i.e. 18.3 – 3.8 kcal/mol at 0°) in MM and 13.4 kcal/mol (i.e. 14.8 – 1.4 kcal/mol at 36°) in QM are brought about by the repulsions between GlcNAc:O1’ carbonyl oxygen and GlcNAc:O3, which also induces unfavourable amide bond non-planarity (Fig S2A). The smaller barrier of 4.8 kcal/mol (i.e. 8.6 – 3.8 kcal/mol at 0°) at 72° for MM and 6.3 kcal/mol (i.e. 7.7 – 1.4 kcal/mol at 36°) at 96° for QM is reduced due to stabilising H-bonds between the GlcNAc *N*-acetyl and the adjacent monosaccharide units (GlcNAc:N2-H⋯Fuc:O2 and reducing Gal:O2-H20⋯GlcNAc:O1’; Fig. S2B). For a backward rotation (i.e. *anti*-to-*syn*), the barrier is higher – 8.6 or 7.7 kcal/mol in MM or QM, respectively, which suggests why we could not observe any such transition in 1.4 μs MD.

The β dihedral angle stayed *trans* in all the α rotational profile optimisations which is not surprising given the high rotational barriers of 19-20 kcal/mol calculated from NMR experiments [21]. We have optimised Le^x^ tetrasaccharide with *anti/trans* and *anti/cis* conformations at the QM level and the latter was higher in energy by 2.6 kcal/mol. The reasons behind this finding are the intrinsic low stability of the β *cis* isomer as found for GlcNAc by NMR [21] and also the lack of stabilising intramolecular H-bonds.

### Molecular dynamics of LecB/Le^x^ complex

To gain insight into the stabilisation of the rare α dihedral in the crystallographic LecB_PA14_/Le^x^ complex, we performed 600 ns MD simulations using the standard force fields for the protein (ff14SB) [31], carbohydrates (GLYCAM06-j) [23], water (TIP3P) [24] and Ca^2+^ ions [32] (Setup 1, Table 2). In line with previous observations [19, 20], the distance between the two calcium ions increased from the crystallographic value of 3.74 Å (Fig. 5A, yellow) to 4.74 Å (average distance after 150 ns, Fig. 5A, black) due to a too high repulsion. A more advanced set of calcium parameters employing the *C_4_* charge-induced dipole term [33] was thereafter used (Setup 2) but with a similar outcome – the average distance after 200 ns was 4.73 Å (Fig. 5A, red). To model the known charge transfer effects [4, 34] implicitly, we used calcium parametrization including effective polarization [35] (Setup 3). The agreement with the experiment was excellent with average values of 3.80±0.15 Å (Fig. 5A, green). Refining further the latter approach, the Ca^2+^-coordinating carboxylates may be scaled for yet improved description [36]. This setup (Setup 4) gave a fair agreement with the crystallographic distance (4.07±0.14 Å; Fig. 5A, blue). Nonbonded parameters for Ca^2+^, K+ and Cl^-^ ions in different MD setups are gathered in Table S3. Partial charges on the Ca^2+^-coordinating carboxylates in different MD setups are gathered in Table S4.

**Table 2.** Summary of the tested MD protocols with different parameterization of calcium ions: 12-6-IOD [32], 12-6-4 [33] or ECCR [35].

| Setup | $\text{Ca}^{2+}$ | Protein | Water |
| --- | --- | --- | --- |
| 1 | 12-6-IOD | ff14SB | TIP3P |
| 2 | 12-6-4 | ff14SB | TIP3P |
| 3 | ECCR | ff14SB | TIP3P |
| 4 | ECCR | ff14SB/scaled carboxylates | TIP3P |
| 5 | ECCR | ff14SB | SPC/E |
| 6 | ECCR | ff14SB | OPC3 |
| 7 | ECCR | ff14SB | TIP4P |

**Figure 5.**
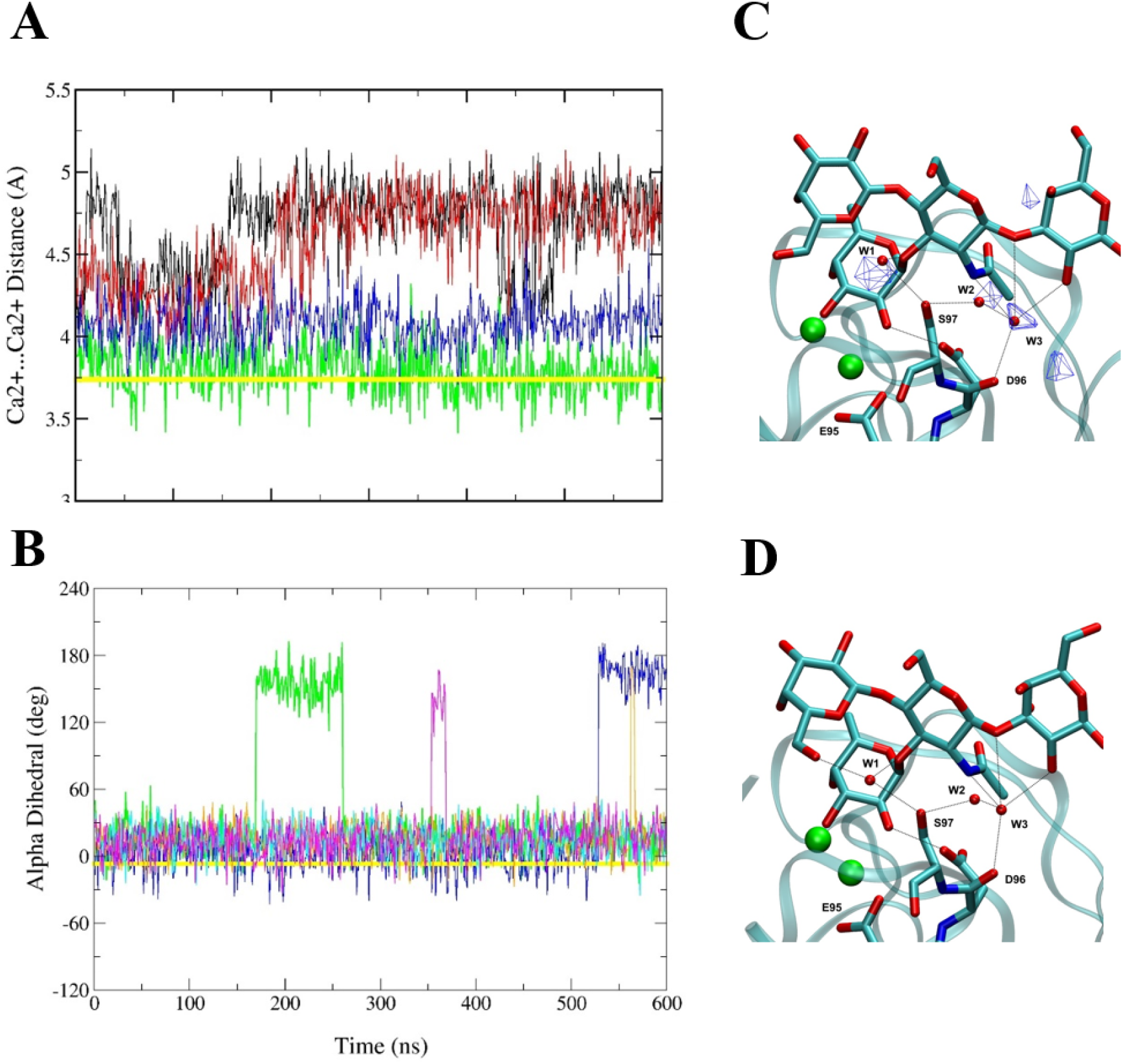
Analyses of 600ns MD simulations of LecB_PA14_/Le^x^ complex and comparison with the crystal structure. **A.** The Ca^2+^⋯Ca^2+^ distance and **B.** the *N*-acetyl α torsion. Curve colour coding: MD simulations with different setups: black – Setup 1; red – Setup 2; green – Setup 3; blue – Setup 4, orange – Setup 5, cyan – Setup 6, magenta – Setup 7. The LecB_PA14_/Le^x^ crystal structure (chain A) determined here – yellow. Zoom at the interaction between LecB (ribbon) and Le^x^ (sticks) from **C.** a representative snapshot from MD using Setup 4 and **D.** the LecB_P_A14/Le^x^ crystal structure. Atom colour coding: cyan – carbon, red – oxygen, blue – nitrogen. Hydrogens are omitted from the MD snapshot for clarity. Dotted lines indicate H-bonding. Blue meshes show densities of water oxygen atoms contoured at 0.06 isosurface value. The densities are 0.112, 0.082 and 0.069 for W1, W2, and W3, respectively.

Further, we analysed the behaviour of the α torsion throughout the MD using Setups 3 and 4, since they maintain the best the architecture of the binding site (as opposed to Setups 1 and 2, c.f. Fig. 5A). Moreover, we tested three other water models in combination with the scaled Ca^2+^ parameters [35] because we knew from the crystal structure that water molecule networks bridge LecB with Le^x^. Besides the classical TIP3P model (Setup 3), these additional water models are: another three-site rigid SPCE model [37] (Setup 5), a more recent three-site OPC3 model [38] (Setup 6) and TIP4P four-site model [39] (Setup 7).

In Fig. 5B (green) we see that MD with Setup 3 maintained the *N*-acetyl group in the α-*syn* conformation only partially (for 85%). After 170 ns, it departed to α-*anti* for 90 ns but thereafter it stayed at α-*syn* until the end of the trajectory. In Setup 4 (Fig. 5B, blue), the α-*syn* conformation was maintained for 88% of the simulation time, i.e. until 530 ns, then it changed to α-*anti* and stayed until the end of the simulation. For the three water models in combination with the scaled-charge Ca^2+^ model, the α-*syn* conformation was retained even for a larger proportion of time (99, 100 and 98%, for Setup 5, 6, and 7, respectively).

A more detailed insight into the reliability of the computational description of the LecB_PA14_/Le^x^ complex is obtained by monitoring two distances. For the interactions of the *N*-acetyl, it is the D96:OD⋯GlcNAc:N distance (**d1**) which in the LecB_PA14_/Le^x^ crystal structure is 3.7 Å in chain A (c.f. Table 1), i.e. it slightly exceeds the criterion for H-bond formation. For the role of Ser97 in organizing the W1-W3 water network, we monitor the Ser97:OG⋯D96:OD distance (**d2**) which in the crystal structure shows no signs of hydrogen bonding (4.6 Å). In all the MD protocols tested, **d1** varies between 2.8 and 4.5 Å but averages to 3.2 Å (Fig. S3), i.e. forming a direct hydrogen bond, not observed in the LecB_PA14_/Le^x^ crystal structure. We note however that i) a direct H-bond has also been observed in the closely related LecB_PA14_/Le^x^ crystal structure and ii) that even the direct H-bonding did not prevent transient formation of bifurcated H-bond toward W3 site, which in turn mediated the bridge to D96:O (c.f. Fig. 2). The **d2** distance evolved during MD towards a non-native strong H-bond (distance of 2.6 Å), except Setups 4 and 7. In Setup 7, a strong Ser97:OG⋯D96:OD H-bond (2.6 Å) was formed for the starting 400 ns and then was lost. But Ser97 then bulged out to the solvent to a non-native position. In Setup 4, Ser97:OG is not attracted so much by D96:OD, due to the reduced charges on the carboxylates and thus does not make the non-native H-bond. This results in an excellent reproduction of the details of the carbohydrate binding site including the location of the Ca^2+^ ions and water networks in Setup 4 as compared to the crystal structure (Fig. 5C, D). Taken together, the charge-scaling procedure on the calcium ions and their coordinating carboxylates (Setup 4, ECCR2) provides the most balanced description of the dynamics in the LecB/Le^x^ complex.

### Discussion and Conclusions

The properties of isolated GlcNAc have been previously studied by a combination of experimental (NMR) and computational (QM, MD) techniques, which yielded a molecular-level understanding of the structures, dynamics and energetics of several conformational families. Regarding the *N*-acetyl group in GlcNAc, the global minimum of the α dihedral is predicted to be *anti* (161° and 180° for the α and β anomers, respectively) [40]. The shift in the minimum between the anomers is caused by intramolecular hydrogen bonding. In the β anomer, a minor population (13%) of α-syn conformation was predicted from a 5 ns MD simulation [40]. We note, however, that this percentage is dependent both, on the force-field type and version and on the length of simulation, all variables that have been improved since [41]. The β amide bond dihedral was found by NMR to be predominantly in *trans* (98%) with a high barrier of 19-20 kcal/mol to conversion to β *cis* [21].

In oligosaccharides containing GlcNAc, especially those where it serves as a branching point, the location and relative stabilities of the minima of the α dihedral along with the transition barriers may change due to the molecular surroundings. Indeed, we found that a QM rotational profile of Le^x^ tetrasaccharide was governed by repulsive oxygen/oxygen contacts on one hand and stabilising O-H⋯O hydrogen bonds, thus shaping the potential energy curve. The GLYCAM06 carbohydrate force field performed qualitatively well in locating the minima but overestimated their energy difference. This translated into the instability of the α-*syn* dihedral in a 1.4 μs-long MD of Le^x^ in explicit solvent, in agreement with expected lower energy barriers in explicit solvent. Additional relaxation of the system may be due to the pyranose ring chair pucker transitions from the most stable *^4^C_1_* chair, which however occurs rarely [14, 42, 43].

A PDB-wide search showed that GlcNAc-containing oligosaccharides bound to proteins exhibited one major (*anti/trans*) and two minor (*syn/trans* and *anti/cis*) populations (c.f. Fig. 3), for which we inspected the experimental electron densities and found that only 16 *syn/trans* cases (including 8 determined in this work, i.e. 4% of the torsions studied) could be confirmed unequivocally. In contrast, for the β dihedral, in the six cases with good fit in the electron densities we could not confirm the minor *anti/cis* population because the *sp^2^* substituents could both be fitted upon 180° rotation. Our two crystal structures of LecB/Le^x^ complexes showed that the geometry and characteristics of the LecB binding site were sufficient to stabilise the rare α-*syn* conformation of *N*-acetyl of GlcNAc. The free energy cost associated with this rearrangement was partly compensated for by acquired hydrogen bonding (direct or water bridged) between the NH group of the *N*-acetyl and D96, as could be observed in the crystal structures.

The LecB carbohydrate-binding site presents a particularly challenging system for force-field calculations due to the charge delocalisation induced by the two closely positioned calcium ions [4]. Previous modelling studies of LecB complexed with monosaccharides indicated problems arising from modelling this special site using the standard 2+ charge model of calcium ion - the Ca^2+^⋯Ca^2+^ distance rose by up to 20 % with respect to the crystallographic one. [19, 20]. Furthermore, the present structures of LecB/Le^x^ complexes determined add another layer of complexity due to the rare α torsion of GlcNAc, whose force-field parametrisation needs to be validated against QM data. Lastly, the presence of water networks mediating protein/carbohydrate interactions requires testing of several water models. A new paradigm in modelling Ca^2+^ ions via effective polarisation had been proposed [44] and developed for inorganic salts, proteins and lipids [35, 36, 45]. The strategy is to scale calcium ion (and counterion) charges by 0.75, reparametrize Lennard-Jones parameters and optionally scale the charged groups in biomolecules. Herein, both these approaches applied to the LecB/Le^x^ complex yielded results superior to standard treatment of calcium ions with 2+ partial charge. Upon testing three additional water models (SPCE, OPC3 and TIP4P) coupled with scaling Ca^2+^ and counterions only, we obtained a better agreement with the crystallographic finding. Finally, we utilised scaled-charge Ca^2+^ ions and counterions which were coupled with TIP4P water model. This setup gave an excellent agreement with the experimental Ca^2+^⋯Ca^2+^ distance and an excellent preservation of the crystallographic α-*syn* dihedral of Le^x^ although with appearance of new hydrogen bonds. The second promising setup (Setup 4) utilised scaled-charge Ca^2+^ ions, counterions and Ca^2+^-coordinating carboxylates and the system was immersed in TIP3P water model. Such an approach gave poorer but still acceptable Ca^2+^⋯Ca^2+^ distance but less preservation of the rare unstable α-syn dihedral of Le^x^. On the contrary, due to the scaling of the binding site carboxylates, occurrence of non-native H-bonding is avoided.

In conclusion, to the best of our knowledge, this is the first report on comprehensive testing of force-field parametrisations on protein with two closely positioned calcium ions in the binding site complexed with a carbohydrate in a rare conformation. By identifying one promising setup, we pave the way for further development, exploration and fine-tuning of parameters to obtain a general protocol for reliable description of such difficult systems where quantum phenomena play important roles. Even though further improvements of force fields are needed for a balanced handling of charge delocalization, the present study allowed for a better understanding of protein/carbohydrate interactions, which can in turn be applied for design of glycomimetics of therapeutical interest.

## Methods

### X-ray crystallography

Molecular cloning, expression and purification of LecB_PA14_ and LecB_PAO1_ were performed as previously described [4, 6]. For crystallization, either LecB_PA14_ or LecB_PAO1_ was dissolved in water (10 mg mL^-1^) and incubated with 800 μg.mL^-1^ of Le^x^ tetrasaccharide (Elicityl) supplemented with 2 mM CaCl2 for 30 minutes prior to crystallization experiment. Crystallization was performed by the hanging drop vapour diffusion method using 1 μL of the protein solution with Le^x^ and 1 μL of the reservoir solution containing 28%-30% PEG8K,

2 mM CaCl_2_, 0.2 M ammonium sulphate, and 0.1 M Tris-HCl pH 8.5 at 19 °C in a 24 well plate. Crystals were cryo-protected with well solution supplemented with 10-25% (v/v) glycerol and flash-cooled in liquid nitrogen. Data were collected at ESRF-BM30A (Grenoble, France) for LecBPA14 or at SOLEIL-PROXIMA1 (Saint Aubin, France) for LecBPAO1, using an ADSC Q315 CCD detector and a Pilatus 6M hybrid photon counting detector (Dectris), respectively. The recorded data were indexed, integrated and scaled using XDS [46] and merged using AIMLESS [47]. The structures were solved by molecular replacement with PHASER [48] using the PDB entry 5A6Q (for LecB_PA14_-Le^x^) or 1W8H (for LecB_PAO1_-Le^x^) as a searching template. The model was finalised by further iterations of manual rebuilding in COOT [22] and restrained refinement in REFMAC5 [49]. Ligand libraries were created using Ligand Builder in Coot and built by Acedrg in CCP4 suite [50]. The geometries of the final models were validated with MOLPROBITY [51], wwPDB validation service (https://validate-rcsb-1.wwpdb.org/) and PDB-redo (https://pdb-redo.eu/) before submission to the Protein Data Bank. All structural figures were prepared using CCP4MG [52]. Data processing and refinement statistics are provided in Table S1 in the Supplementary Information.

### Protein Data Bank (PDB) Search

The values of the N-acetyl α and β dihedrals of β-D-GlcNAc pyranose residues in protein/carbohydrate complexes were measured in the PDB database using the GlyTorsion tool [53]. To increase the confidence of the structural parameters, we set the X-ray resolution cutoff to 1.5 Å. NMR structures were filtered out for the lack of a quality validation criterion. To be able to observe the effect of the neighbouring carbohydrate moieties, we set the minimal oligosaccharide chain length to trisaccharide. The GlyTorsion tool measures the ω2a (C1-C2-N-C1’) and ω2b (C2-N-C1’-O1’) dihedrals (“x-ray definition”, i.e. using heavy atoms only, c.f. Fig. 1D). These values were transformed into a more readily understandable “NMR definition” [7] (using hydrogens added via LEaP program of AMBER18 [30]: α dihedral (H2-C2-N-H) and β dihedral (H-N-C-O) by adding the approximate value of 60° (deviations of several degrees from the exact values, data not shown) and the exact value of 180°, respectively. This approach avoids the need of adding hydrogens to the structures in a PDB-wide analysis.

### Modelling

All the modelling was done on A chain (or A/B chain dimer) of LecB_PA14_/Le^x^ structure because of the high quality of the electron density maps of the whole Le^x^ tetrasaccharide. Crystallographic water molecules were retained except those with partial occupancy which may induce clashes. Hydrogens were added using the LEaP program of AMBER18 [30]. As suggested by the previous crystal structure and semiempirical quantum chemical calculations [4], the carboxylates in the active site were left unprotonated. The molecular mechanics (MM) force fields used were: ff14SB [31] for the protein and GLYCAM-06j [23] for the carbohydrate. Several Ca^2+^ parametrisations were used: standard nonbonded +2 charge [32] optimized for ion-oxygen distance (IOD) (Setup 1); more advanced one +2 charge and employing the *C_4_* charge-induced dipole term (Setup 2) [33], a setup with charge on calcium and counterions scaled by 0.75 (Setup 3) [35] and lastly a setup inspired by [36] where Setup 3 was augmented with scaling Ca^2+^-coordinating carboxylates by 0.75 (Setup 4; partial charges are in Table S4). Several water models were used in conjunction with the scaled calcium and counterion setup: TIP3P [24] (Setup 3), SPCE [37](Setup 5), OPC3 [38](Setup 6) and TIP4P [39] (Setup 7). Hydrogens were optimized using the generalized Born (GB) implicit solvent model [29] (ε_r_ of 78.5) by 5000 cycles of LBFGS optimization with zeroed dihedral barriers, followed by another 5000 LBFGS cycles in full force field.

### Quantum Chemical Calculations

*N-Acetyl Rotational Scan*. The α dihedral in the Le^x^ tetrasaccharide was rotated to 0° using Cuby4 [54]. Thereafter it was rotated stepwise by 12° in either direction up to a dihedral of - 180/180° and optimised in each step to tight convergence criteria (maximum energy difference of 0.6 cal/mol, maximum gradient of 0.12 kcal/mol/Å^2^), keeping the α dihedral fixed. This was performed in Cuby4 [54] at MM/GB level (using GLYCAM06-j [23] for Le^x^ and *igb=5* option for generalised Born implicit solvation) using AMBER18 [30] or QM/COSMO level (DFT-D3 using B-LYP functional/DZVP basis set [26]) with COSMO [27] implicit solvent using Turbomole 7.0 [28]. Energies of the relaxed structures relative to the minimum of the respective method were plotted.

*QM Charge Calculations*. The LecB_PA14_ chain A/B dimer with two Le^x^ tetrasaccharides, four calcium ions and the crystallographic water molecules had its hydrogen atoms optimized by use of MM/GB [29]. Thereafter, it was subjected to 600 cycles of further hydrogen optimization at the semiempirical QM (SQM) level using PM6-D3H4 [55] coupled with COSMO [27] using MOPAC2016 (http://OpenMOPAC.net). The resulting structure was subjected to another 600 cycles of hydrogen optimization at QM/SQM/COSMO level, QM being B-LYP/DZVP-D3 [26]. We used a subtractive mechanical embedding scheme with link hydrogen atom approach. The QM part consisted of residues within 4.8 Å of the two calcium ions in the A chain, the Le^x^ tetrasaccharide and one crystallographic water molecule (W1), altogether 262 atoms. The SQM part comprised the whole system. For charge calculations, Natural Bond Orbital (NBO) [56] method was used on the isolated QM part or isolated and its QM/COSMO optimized constituents

### Molecular Dynamics

The LecB chain A/B dimer with two Le^x^ tetrasaccharides, four calcium ions and the crystallographic water molecules with MM/GB optimized hydrogens was immersed in an octahedral box of water molecules which extended at least 12 Å from the solute. K^+^/Cl^-^ counterions (scaled in Setups 3 and 4; their Lennard-Jones parameters are shown in Table S3) were added to maintain neutrality and physiologic concentration of 0.15 M. Bonds involving hydrogen atoms were constrained using SHAKE [57]. Hydrogen mass repartitioning to 3Da [58] was applied, which allowed us to use a longer integration time step of 4 fs. Initial stepwise relaxation and subsequent MD was performed according to the published protocol [59].

## ASSOCIATED CONTENT

The crystallographic methodology and analyses, QM optimised structures, MD analyses and MD parameters are presented in the Supporting Information.

## CONTRIBUTORS

M. Lepsik, AI and AT designed the study. RS and SK performed the crystallography work under the guidance of AV. M. Lepsik performed all calculations with the help of M. Lelimousin and EP. M. Lepsik, SK and AI wrote the manuscript.

## ACKNOWLEDGEMENT

Dr. Martin Lepsik has received funding for this project from the European Union’s Horizon 2020 research and innovation programme under the Marie Sklodowska-Curie grant agreement No 795605. The authors acknowledge support by the ANR PIA Glyco@Alps (ANR-15-IDEX-02), Labex ARCANE and CBH-EUR-GS (ANR-17-EURE-0003). Further, we acknowledge funding from the Helmholtz Association (grant no. VH-NG-934). We are grateful to synchrotron SOLEIL (Saint Aubin, France) and ESRF (Grenoble, France) for access and technical support at beamline PROXIMA 1 and BM30A, respectively. Part of the computations presented in this paper were performed using the Froggy platform of the CIMENT infrastructure which is supported by the Rhône-Alpes region (GRANT CPER07_13 CIRA) and the Equip@Meso project (reference ANR-10-EQPX-29-01). The work has been performed under the Project HPC-EUROPA3 (INFRAIA-2016-1-730897), with the support of the EC Research Innovation Action under the H2020 Programme; in particular, the author gratefully acknowledges the computer resources and technical support provided by EPCC at the University of Edinburgh, Scotland. We thank Emilie Gillon for providing recombinantly produced LecB_PAO1_ for X-ray crystallographic experiments.

