## Supporting Information for "Induction of Rare Conformation of Oligosaccharide by Binding to Calcium-dependent Bacterial Lectin: X-ray Crystallography and Modelling Study"

**Table S1**. X-ray data collection and processing of LecB structures

| **Data set** | **LecB_PAO1_­- Le^x^** | **LecB_PA14_- Le^x^** |
| --- | --- | --- |
| **PDB code** | 6R35 | 5A70 |
| **Data Collection** |  |  |
| **Beamline** | PROXIMA1 (SOLEIL) | BM30A (ESRF) |
| **Wavelength (Å)** | 0.9792 | 0.9802 |
| **Detector** | Pilatus 6M | ADSC Q315 CCD |
| **Resolution (Å)^a^** | 47.82-1.84 (1.80-1.80) | 44.94-1.60 (1.63-1.60) |
| **Space Group** | P2_1_ | P2_1_ |
| **a, b, c (Å)** | 52.60, 72.48, 62.06 | 53.0, 63.3, 63.9 |
| **α, β, γ (°)** | 90.0,114.6 ,90.0 | 90.0, 91.2, 90.0 |
| **Total observations** | 203335 | 161897 |
| **Unique reflections** | 39320 | 53856 |
| **Multiplicity^a^** | 5.2 (5.0) | 3.0 (2.3) |
| **Mean *I*/σ(*I*)^a^** | 13.7 (3.1) | 17.5 (2.5) |
| **Completeness (%)^a^** | 99.9 (99.6) | 96.8 (74.7) |
| ***R*_merge_^a,b^** | 0.076 (0.533) | 0.041 (0.330) |
| ***CC*_½_^a,c^** | 1.0 (0.9) | 1.0 (0.8) |
| **Refinement** |  |  |
| **Reflections: working/free^d^** | 39301/2004 | 50981/2857 |
| ***R*_work_/ *R*_free_^e^** | 0.132/0.176 | 0.136/0.171 |
| **Ramachandran plot: allowed/favoured/outliers (%)^f^** | 100/97.8/0 | 100/97.5/0 |
| **R.m.s. bond deviations (Å)** | 0.0172 | 0.0170 |
| **R.m.s. angle deviations (°)** | 1.869 | 1.700 |
| **Mean *B*-factors: protein/ligand^f^/**  **/water (Å^2^)** | 19/36/31 | 13/20/27 |

^a^ Values for the outer resolution shell are given in parentheses.

^b^ *R*_merge_ = ∑_hkl_ ∑_i_ |I_i_(hkl) − 〈I(hkl)〉|/ ∑_hkl_ ∑_i_I_i_(hkl).

^c^ *CC*_½_ is the correlation coefficient between symmetry-related intensities taken from random halves of the dataset.

^d^ The data set was split into "working" and "free" sets consisting of 95 and 5% of the data, respectively. The free set was not used for refinement.

^e^ The R-factors *R*_work_ and *R*_free_ are calculated as follows: *R* = ∑(| *F*_obs_ - *F*_calc_ |)/∑| *F*_obs_ |, where *F*_obs_ and *F*_calc_ are the observed and calculated structure factor amplitudes, respectively

^f^ refers to ligands bound in the active site and potential surface binding sites

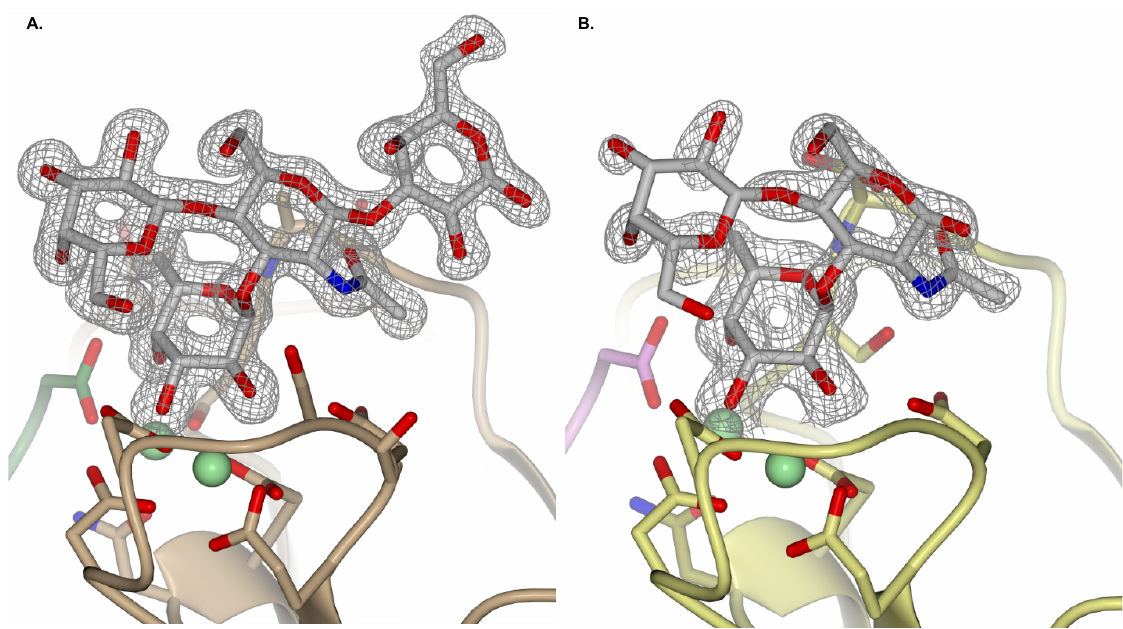

**Figure S1.** Electron density map for Le^x^ in complex with two LecB variants, **A.** LecB_PA14_ and **B.** LecB_PAO1._ The electron density is displayed at 1σ. Figure was prepared with CCP4MG.

**Table S2.** Φ, Ψ torsion angles (°) for the two glycosidic linkages of Le^x^ trisaccharide core in the crystal structures of LecB_PA14_/Le^x^ and LecB_PAO1_/Le^x^ complexes.

|  | LecB_PA14_/Le^x^ | | | | LecB_PAO1_/Le^x^ | | | |
| --- | --- | --- | --- | --- | --- | --- | --- | --- |
|  | *α*Fuc1-3GlcNAc | | *β*Gal1-4GlcNAc | | *α*Fuc1-3GlcNAc | | *β*Gal1-4GlcNAc | |
| Chain | Φ _O5-C1-O1-C3_ | Ψ _C1-O1-C3-C4_ | Φ _O5-C1-O1-C4_ | Ψ _C1-O1-C4-C5_ | Φ _O5-C1-O1-C3_ | Ψ _C1-O1-C3-C4_ | Φ _O5-C1-O1-C4_ | Ψ _C1-O1-C4-C5_ |
| A | -69.6 | 136.9 | -72.6 | -103.8 | -86.1 | 146.6 | -67.6 | -107.8 |
| B | -76.8 | 140.7 | -68.1 | -108.7 | -87.4 | 148.1 | -58.4 | -110.9 |
| C | -70.9 | 139.2 | -72.2 | -107.4 | -83.8 | 150.7 | -63.0 | -104.3 |
| D | -71.4 | 138.4 | -68.3 | -107.7 | -70.5 | 138.9 | -73.3 | -107.8 |

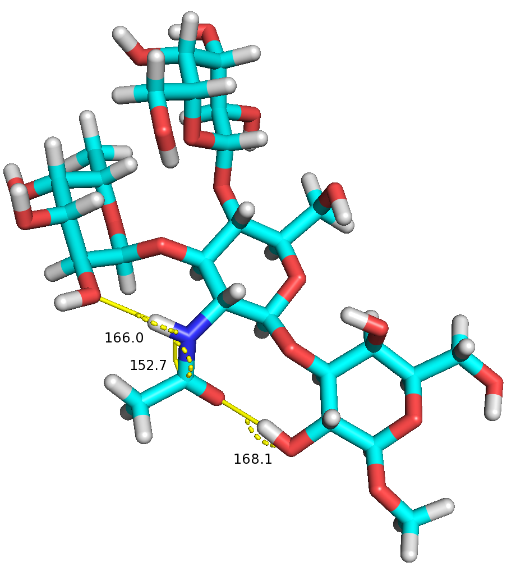

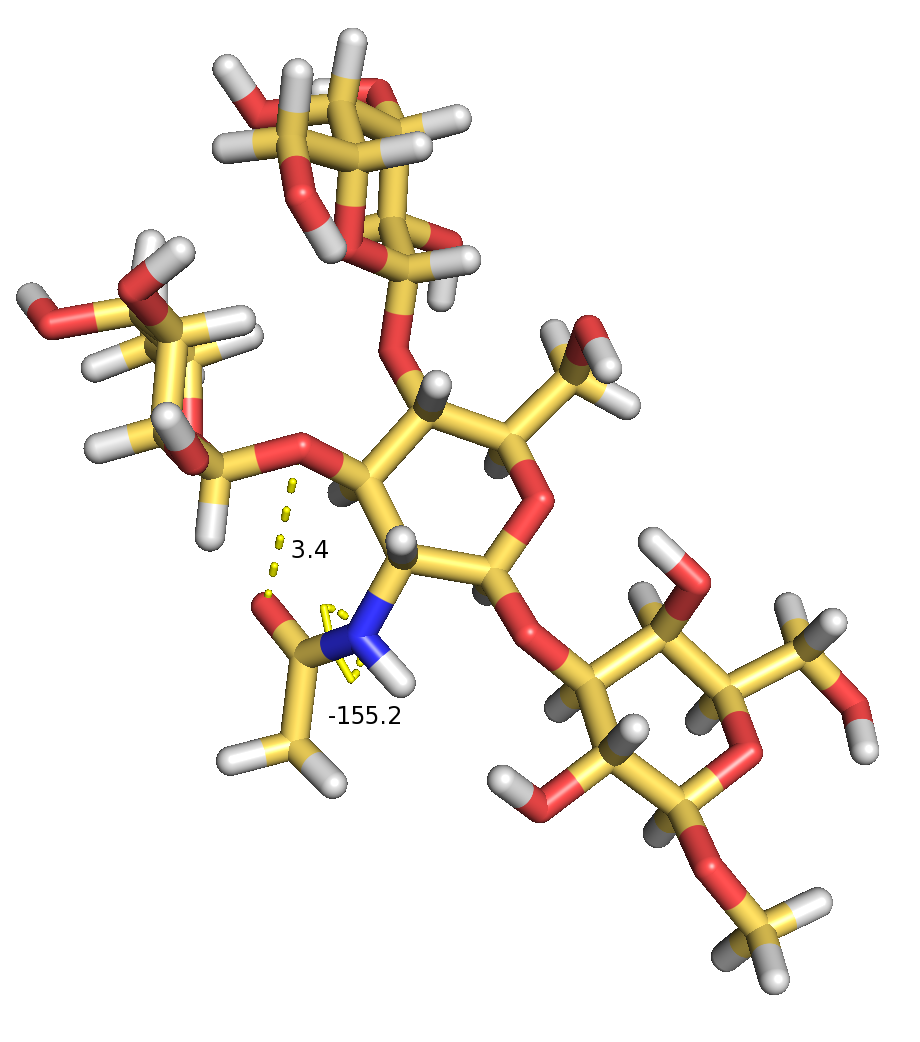

**Figure S2.** Optimised structures of Le^x^ from rotational profile at QM/COSMO level. **A.** Barrier at -96**°**, **B.** Barrier at 96°. Colour coding: red – oxygen, blue – nitrogen, white hydrogen, yellow/cyan – carbon. Distances are in Å, dihedral angles in degrees. Figure prepared with PyMol, version 1.8.

**Table S3.** Nonbonded parameters for Ca^2+^, K^+^, Cl^-^ ions in different MD setups

| Setup | 1 | 2 | 3 | 4 |
| --- | --- | --- | --- | --- |
|  | 12-6 IOD | 12-6-4 | ECCR | ECCR2 |
| Ca^2+^ |  |  |  |  |
| charge (e) | 2.0 | 2.0 | 1.5 | 1.5 |
| σ (GROMACS, Å) |  |  | 2.5376 | 2.6656 |
| R_min_/2 (AMBER, Å) | 1.6080 | 1.6420 | 1.4236 | 1.4954 |
| ε (GROMACS, kJ/mol) | |  | 0.5072 | 0.5072 |
| ε (AMBER, kcal/mol) | 0.0830 | 0.1019 | 0.1212 | 0.1212 |
| Cl^-^ |  |  |  |  |
| charge (e) | -1.00 | -1.00 | -0.75 | -0.75 |
| σ (GROMACS, Å) |  |  | 3.7824 | 4.1000 |
| R_min_/2 (AMBER, Å) | 2.1620 | 2.1500 | 2.1219 | 2.3001 |
| ε (GROMACS, kJ/mol) |  |  | 0.4184 | 0.4928 |
| ε (AMBER, kcal/mol) | 0.5315 | 0.5215 | 0.1000 | 0.1178 |
| K^+^ |  |  |  |  |
| charge (e) | 1.00 | 1.00 | 0.75 | n.d |
| σ (GROMACS, Å) |  |  | 3.154 | n.d |
| R_min_/2 (AMBER, Å) | 1.7450 | 1.7580 | 1.7694 | n.d |
| ε (GROMACS, kJ/mol) |  |  | 0.4187 | n.d |
| ε (AMBER, kcal/mol) | 0.1702 | 0.1800 | 0.1001 | n.d |

**Table S4.** Partial charges on the Ca^2+^-coordinating carboxylates in different MD setups

|  |  | Setups 1-3 | Setup 4 |
| --- | --- | --- | --- |
| ASP | OD1 | -0.8014 | -0.6011 |
|  | CG | 0.7994 | 0.5996 |
| GLU | OE1 | -0.8188 | -0.6141 |
|  | CD | 0.8054 | 0.6041 |
| G114* | O | -0.7855 | -0.5891 |
|  | C | 0.7231 | 0.5423 |

* C-term of B-chain residue

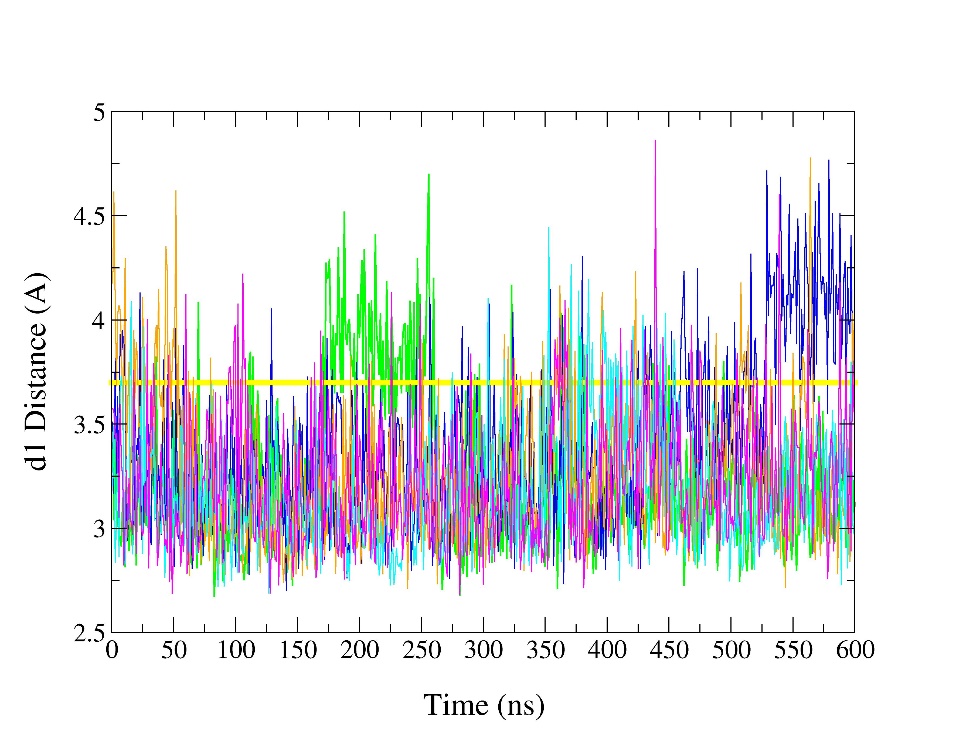

**Figure S3.** Evolution of the **d1** distance in the tested MD protocols and comparison with the LecB_PA14_/Le^x^ crystal structure (chain A) determined here - yellow, MD colour coding: black – Setup 1; red – Setup 2; green – Setup 3; blue – Setup 4, orange – Setup 5, cyan – Setup 6, magenta – Setup 7.
